# Effects of dietary NFC/NDF on rumen microbiomes of Karakul sheep based on Three Generations of Full-length Amplifiers sequencing

**DOI:** 10.1101/729780

**Authors:** Xuanxuan Pu, Xuefeng Guo, Chenyu Jiang, Junfeng Liu, Xiuping Zhang, Sujiang Zhang, Long Cheng, Anshan Shan

## Abstract

An study was was conducted to investigate the effects of dietary(non fibrous carbohydrate) NFC/(neutral detergent fiber)NDF on ruminal bacteria in Karakul sheep. Twelve Karakul sheep were assigned randomly to four dietary treatments of NFC/NDF (0.78, 1.23, 1.61 and 2.00 respectively) as group 1, 2, 3 to 4. The experiment lasted for four periods, period I (1~18 d), II (19~36 d), III (37~54 d) and IV (55~72 d). Ruminal digesta were collected consecutively for three days to measure pH and bacteria per period. The results indicated that the average ruminal pH and amounts of OTUs were decreased with the increase of dietary NFC/NDF for four periods. At phylum level, Bacteroidetes and Firmicutes were the predominant bacteria of four periods, Bacteroidetes were decreased, while the relative abundance of Firmicutes was increased with dietary NFC/NDF for four periods, but the difference wasn’t significant (*P*>0.05). At genus level, the most relative abundance genus was unidentified-Lachnospiraceae which reached the highest in group 3 for four periods, but the difference wasn’t significant (*P*>0.05). Conclusion: ruminal pH and bacteria were decreased with the increase of dietary NFC/NDF and the most dominant bacteria were not change with dietary NFC/NDF and periods in Karakul sheep.

## Introduction

Rumen plays an important role in the growth and production of ruminants and it contains a large number of rumen microorganisms (*Bacteria*, *Protozoa*, *Eukarya* and *Archae*a[1–3]). Rumen microorganisms breakdown food to provide volatile fatty acids, bacterial protein, and so on to the host animal[4]. Yang et al.[5] confirmed that 80% of the starch, 50% of the fibre and 60% of the organic matter in the diet were fermented in the rumen to provide energy for the host. Meanwhile, after a long-term selection and evolution, rumen microbiomes and the host have formed a symbiotic relationship to maintain the host’s health[6]. Dietary regulation has an important effect on rumen fermentation[7–9] and microbiome[10–11]. Wei et al.[12] showed that with the increase of dietary NFC/NDF, ruminal pH decreased significantly, furthermore the composition of rumen microbial flora also changed, and the total number of rumen bacteria decreased in goats. However, there are scarce studies on rumen microbiome structure changes with dietary NFC/NDF in Karakul sheep and the rumen is very complex in which microbiome may change again with prolong of periods. NDF plays an important role in dry matter intake (DMI) and feed digestibility[13–14], and NFC in diets is another factor that affect DMI, Hall et al. reported that NFC would be degraded rapidly in the rumen[14]. The development of technology makes it more accurate to study on rumen microbiomes, Three Generation of Full-length Amplifiers sequencing is one of them, which can improve the resolution of species identification, and improve the accuracy of microbial species composition identification in the samples[15–16]. In this experiment, ruminal pH and microbiome were measured for four periods to investigate effects of dietary NFC/NDF on ruminal microbiome in Karakul sheep.

## Materials and Methods

### Animals and Dietary composition

All experimental procedures were approved by Tarim University Animal Care and Use Committee, and humane animal care were followed throughout the experiment. Twelve Karakul sheep with similar age and weight (35.3 ± 3.3 kg) were fitted with permanent fistula and were randomly assigned into four dietary treatments of NFC/NDF (0.78, 1.23, 1.61, 2.00 respectively) as group 1, 2, 3 and group 4, each group with three replicates. They all received vaccines for parasites before the adaption period, and were fed meeting the standards for raising meat and sheep in the People’s Republic of China[17]. All sheep were housed individually in metabolic cages (1.2 m × 1.5 m) and fed the experimental diet individually twice a day at 9:00 a.m. and 8:00 p.m with free access to water. The ingredients and nutrient level of the diet were shown in S1 Table.

### The experimental design and sample collection

The experiment lasted for 72 d including four periods, including period I (1~18 d), II (19~36 d), III (37~54 d) and IV (55~72 d). Each period lasted for 18 d. The first 15 d for adaption and 3 d for samples, feed intake and defecation rule were studied in the adaption period. The ruminal digesta were sampled consecutively before morning feeding for three days, the sheep of one group were collected together for 50 mL, and then, were stored in −80 ℃ to investigate on rumen microorganisms. Meanwhile, the ruminal pH was measured after feeding of 0, 1, 3, 6 and 9 h using pH meter (FE22).

### DNA Extraction, PCR and Pacio sequencing

The total genetic DNA was extracted using QIAamp Fast DNA Stool Mini Kit (QIAGEN, Shanghai) according to the illustration and the extracted DNA was detected by 1% agarose gel. The V1-V9 regions of 16S rDNA were amplified by PCR from the extracted DNA using the universal primers: F, 5’-AGAGTTTGATCCTGGCTCAG-3’; R, 5’-GNTACCTTGTTACGACTT-3’ (synthesized by Biological engineering co., Ltd). PCR was carried out in triplicate 50-μL reactions which containing 2 μL Primer Mix (1uM), 5 ng gDNA, 1 μL Trans Fastpfu, 10 μL 5× Buffer, 5 μL 5× StimuLate, 5 μL dNTPs (2.5mM each), 27 μL NFW. Thermocycling parameters were as follows: 2 min predenaturation at 95 °C; 35 cycles of denaturation at 95 ℃ for 30 s, Annealing at 60 ℃ for 40 s, extension at 72 ℃ for 90 s; and a Final extension at 72 ℃ for 10 min. The production was detected by 2% agarose gel. PCR products was purified with Gel Extraction Kit (QIAGEN, Shanghai), and the productions were sequenced on PacBio platform.

### The sequence analysis

The 16S rDNA reads were firstly processed to get clean reads by discarding the reads that are shorter than 1340 bp, longer than 1640 bp, and not matching the expected barcodes. Using Uparse software [18] to cluster all Clean Reads of all samples. Operational taxonomic units (OUTs) were formed at the similarity of 97%[19]. The OUTs were annotated by the Mothur and SILVA (http://www.arb-silva.de/) [20] according to the reference taxonomy provided by SSUrRNA detabase[21]. The OTUs were analyzed by Qiime pipeline (Version 1.9.1) to calculate the richness and diversity indices i e. observed OTUs, Chao1, Shannon, Simpson, ACE.

### Statistical Analyses

The results of pH were expressed as means using SPSS 17.0. Comparisons between groups were performed with ANOVA followed by Duncan test. The difference of ruminal bacteria was measured and expressed using SPSS 17.0. As well, and the differences were considered to be significant at *P*<0.05.

## Results

### pH

The average pH of 0, 1, 3, 6, 9 h after feeding for four periods were shown in Table 1, the pH was that: group 1> group 2> group 3> group 4 for four periods. There was no significant difference between four groups in period I (*P*>0.05), while there was significant difference between four groups in period II, III and IV (*P*<0.05), which showed that the ruminal pH decreased significantly as the dietary NFC/NDF increased.

**Table 1.** Effects of different NFC/NDF on rumen fluid pH

| Items | pH |  |  |  | SEM | *P*-value |
| --- | --- | --- | --- | --- | --- | --- |
|  | 1 | 2 | 3 | 4 |  |  |
| Period I | 6.36 | 6.32 | 6.26 | 6.05 | 0.07 | 0.507 |
| Period II | 6.42a | 6.28ab | 6.03bc | 6.01c | 0.06 | 0.015 |
| Period III | 6.68a | 6.48ab | 6.34bc | 6.19c | 0.06 | 0.012 |
| Period IV | 6.52a | 6.27ab | 6.16b | 6.05b | 0.06 | 0.016 |

Note: In the same row, values with no letter or the same letter superscripts mean no significant difference ( $P>0.05$ ), while with different small letter superscripts mean significant difference ( $P$ $<0.05$ ). Period I (1~18 d), II (19~36 d), III (37~54 d) and IV (55~72 d) and Group 1, 2, 3, 4 means four groups of sheep treated with four dietary levels of NFC/NDF (0.78, 1.23, 1.61, 2.00 respectively). The same as Table 3.

### Extraction DNA of rumen bacteria

The DNA extraction results were shown in Fig. 1. The main band was clear and there was no concentrated band below 500 bp, which indicated that the purity of DNA was well and it could meet the requirement of sequencing.

**Fig. 1.**
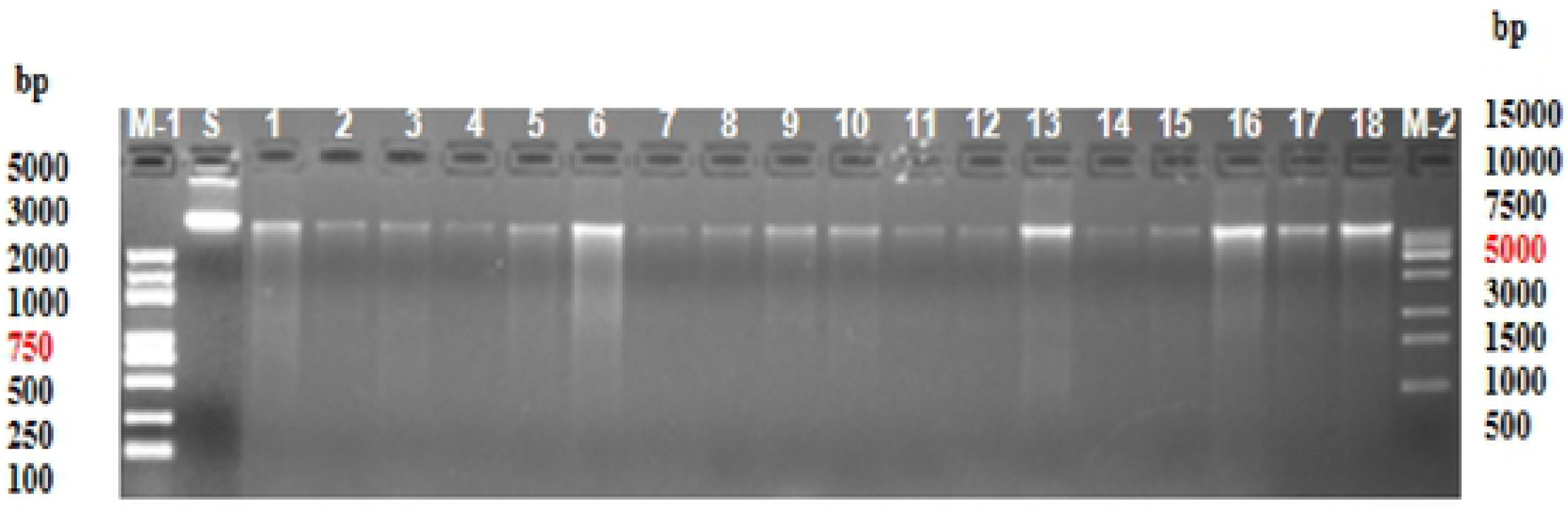
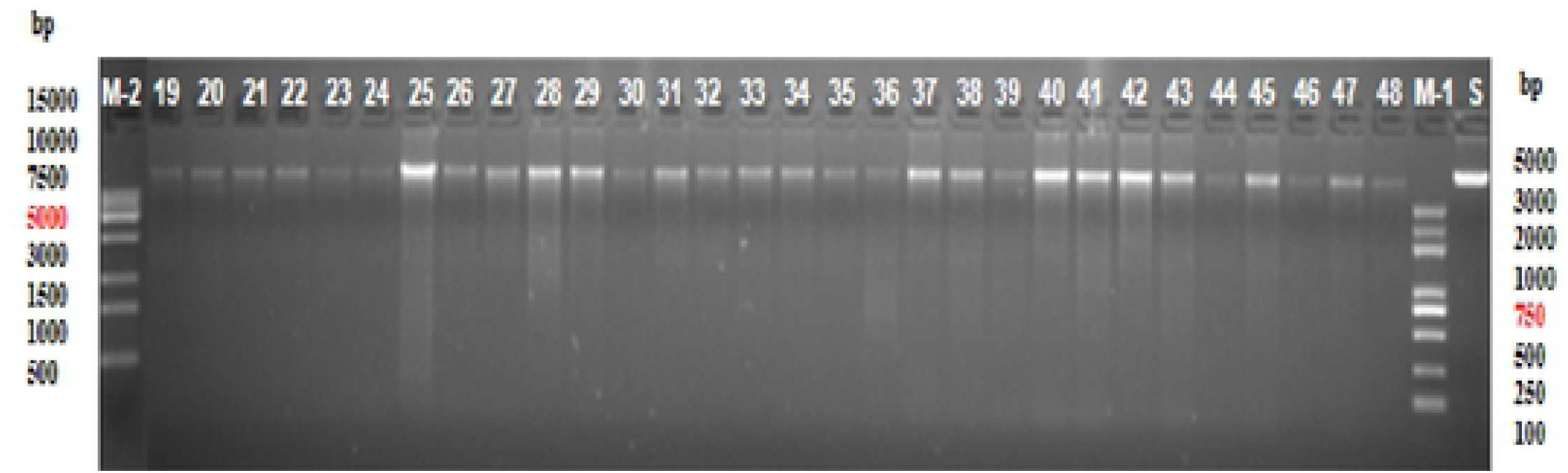
Extraction of rumen bacterial DNA. The DNA of forty-eight samples from four periods samples were extracted, the order of 1~3, 4~6, 7~9 and 10~12 means samples in group 1, 2, 3 and group 4 of period 1 respectively, each group with three replicates. The following order are as period I.

### The analysis of basic sequencing data

The OTU number of each group in one period was shown in Venn graph as Fig. 2. The result showed that the OTUs was that: group 1> group 2 > group 3> group 4 for four periods, which showed that the diversity of rumen microbiome decreased with the increase of NFC/NDF, and the number of each group became more stable with prolong of periods

**Fig. 2.**
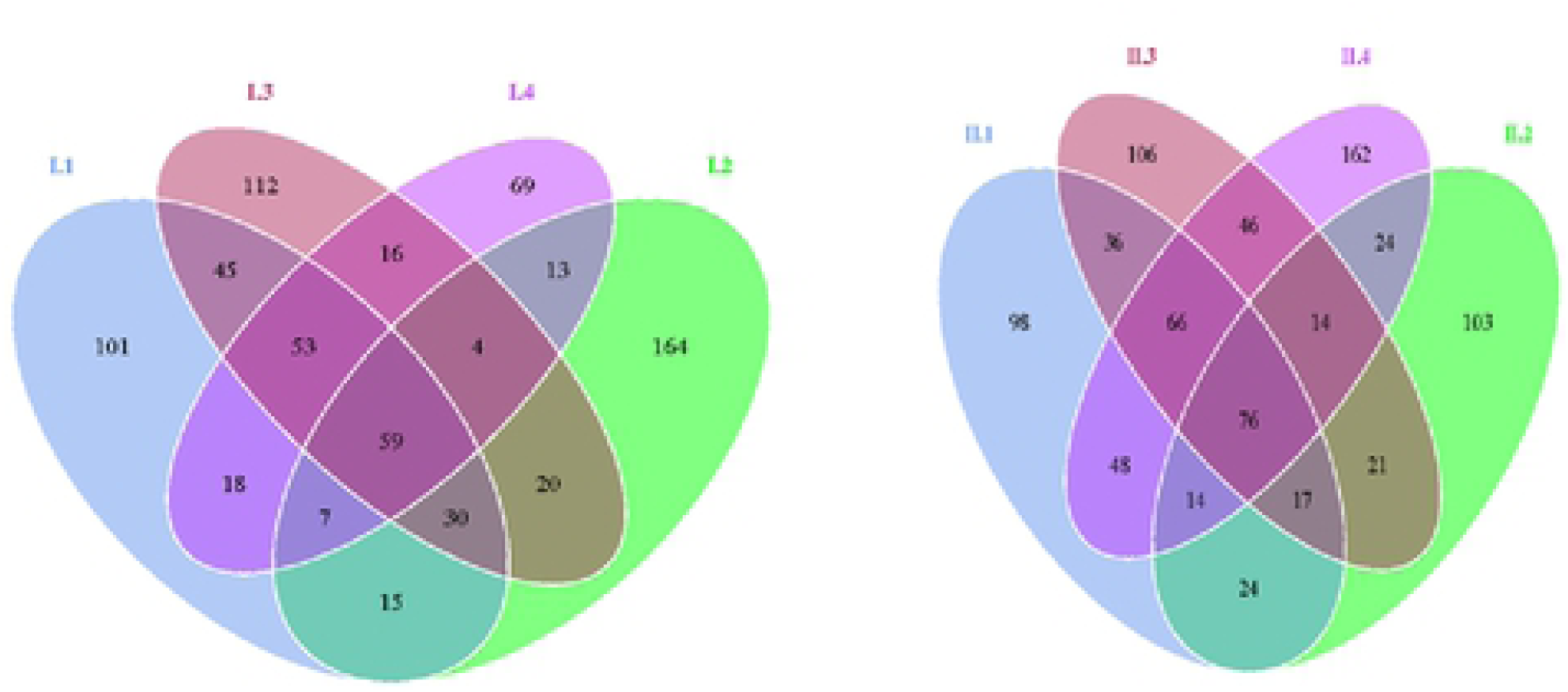
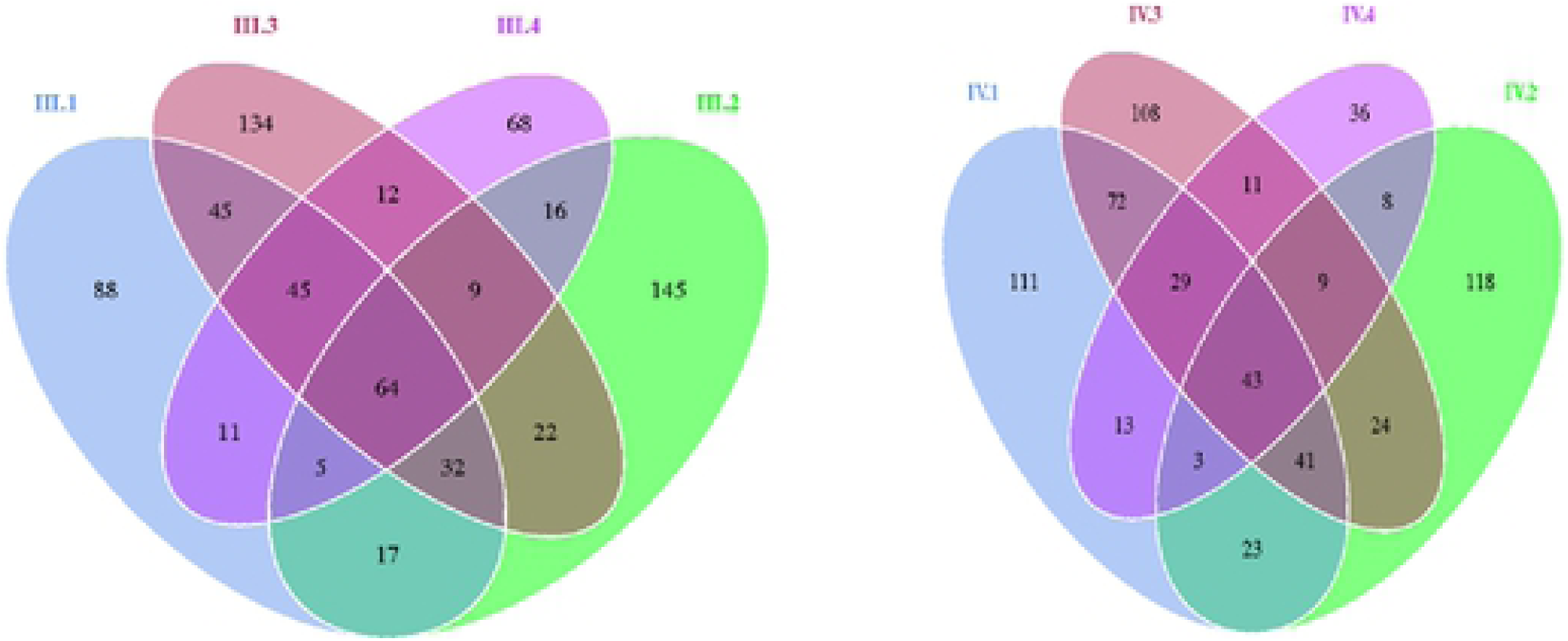
Vene graph of microflora in rumen fluid of Karakul sheep. The amounts of OTUs in each group were shown and four groups of one period were formed in one Vene graph.

### The analysis of OTU Alpha diversity

The Alpha diversity is a kind of analysis in the diversity of microbiome, which involves the abundance index of Chao1[22] and ACE[23] and the diversity index of Shannon and Simpson[24]. Before diversity analysis, the dilution curve was drawn by R software (Version 2.15.3) to detect whether the obtained data could fully reflect the distribution of rumen fluid flora in Karakul sheep.

### OTU dilution curve

As showed in Fig. 3, the dilution curve of each group keep increasing with the increase of the depth of sequencing, which is indicated that new bacteria had been found. The results showed that the sequencing quantity of each sample could be used to analyze the diversity of flora.

**Fig. 3.**
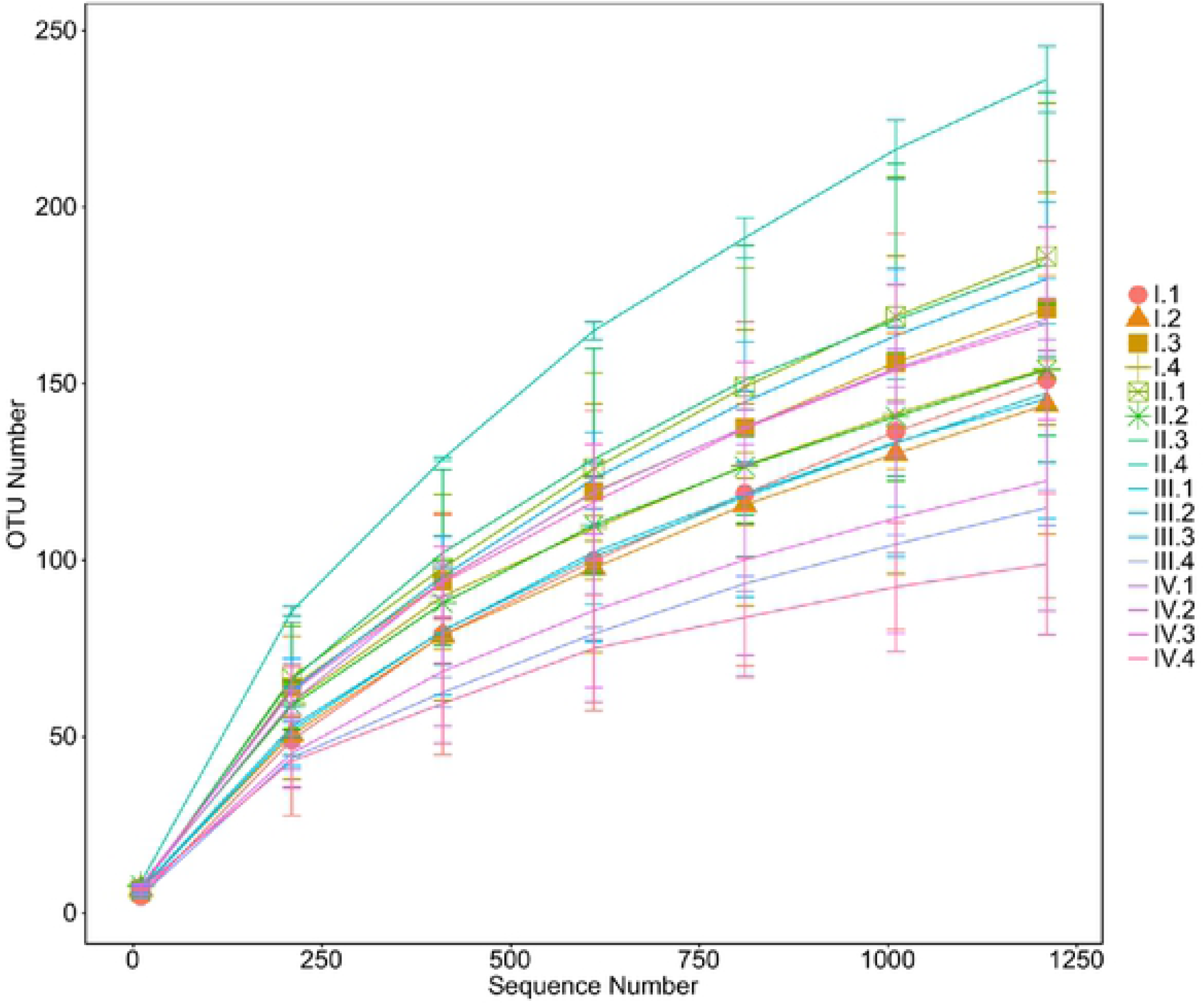
OTU dilution curve of bacteria in the rumen of Karakul sheep. Rarefaction curves of OTUs clustered at 97% sequence identity across different samples.

### Sample diversity index

The results of Alpha diversity were shown in Table 2. The results showed that there was some difference in the number, richness and diversity of rumen bacterial species in different dietary of NFC/NDF.

**Table 2.**
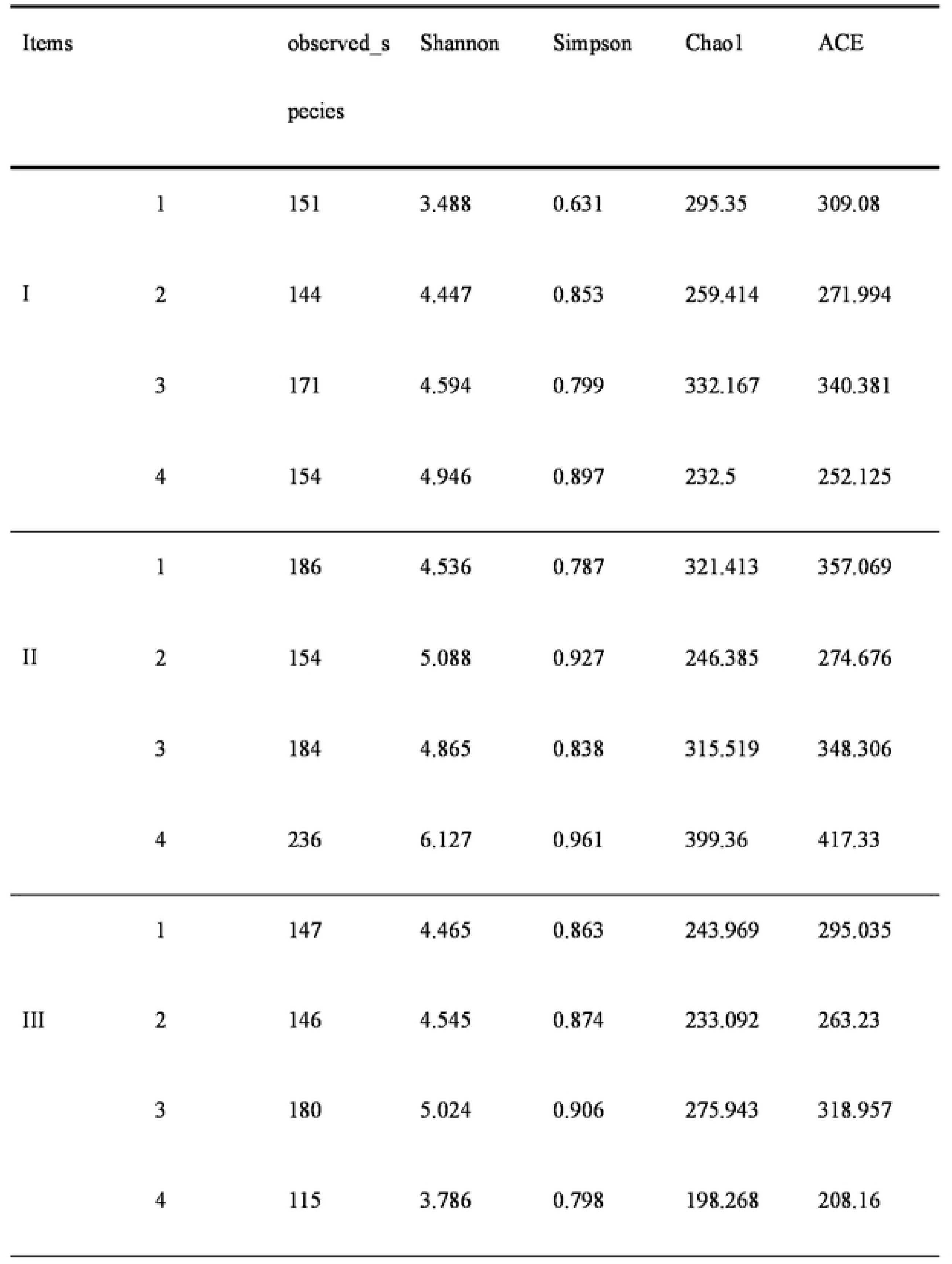
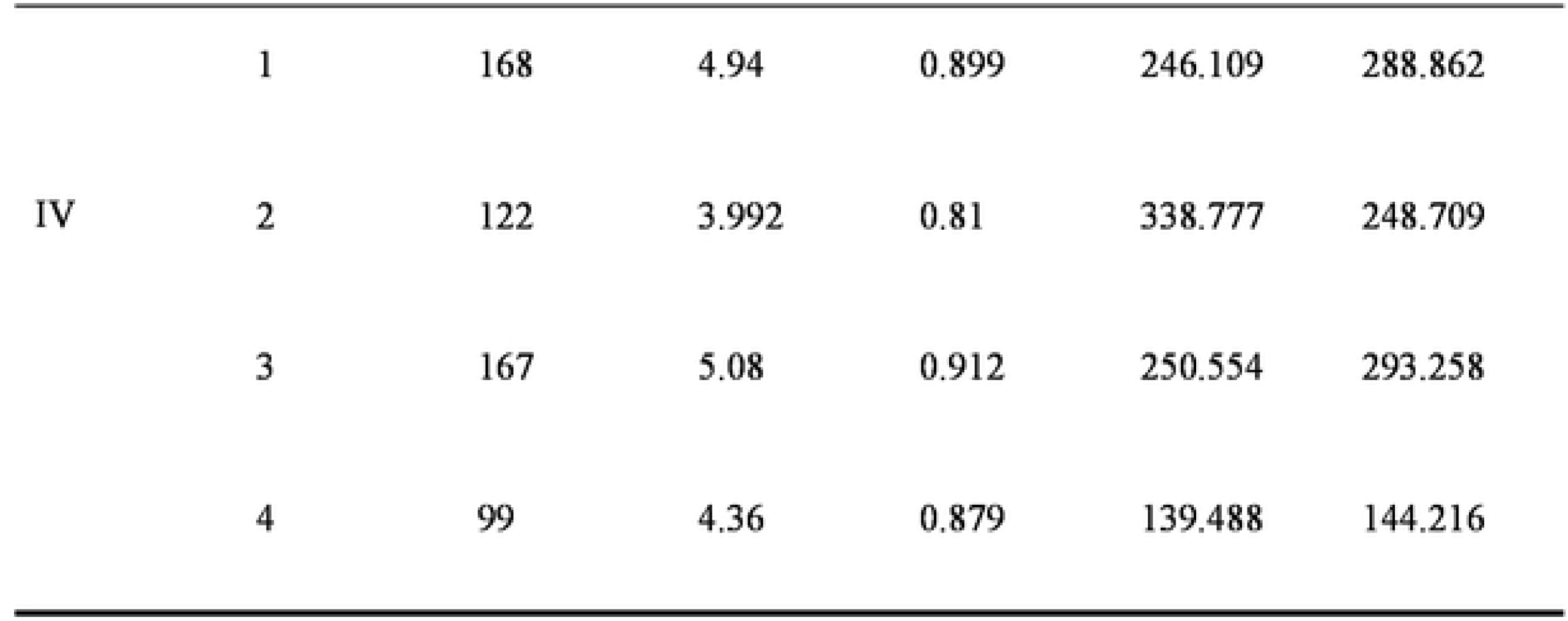
Analysis of Alpha diversity of rumen liquid samples at 0.03 distance

### Structural analysis of rumen microbiome

At phyla level, seventeen different phyla were detected. The S2 table and Fig. 4 showed that the main phylum microbiome didn’t change with the dietary NFC/NDF and periods, and Bacteroidetes (52%~72%) and Firmicutes (19%~46%) was the main microbiome for four periods. The relative abundance of Tenericutes was 3% to 8% and the others was less than 1%. Fig. 5 was composed by four main dominant phylum to investigate effects of NFC/NDF on their relative abundance, the abundance of Bacteroidets and Proteobacteria were that: group 1> group 2> group 3> group 4 for four periods, but the difference wasn’t significant(*P*>0.05). While the abundance of Firmicutes was that : group 4> group 3> group 2> group 1, the difference wasn’t significant(*P*>0.05) as well. The abundance of Tenericutes reached the highest in group 4 for four periods.

**Fig. 4.**
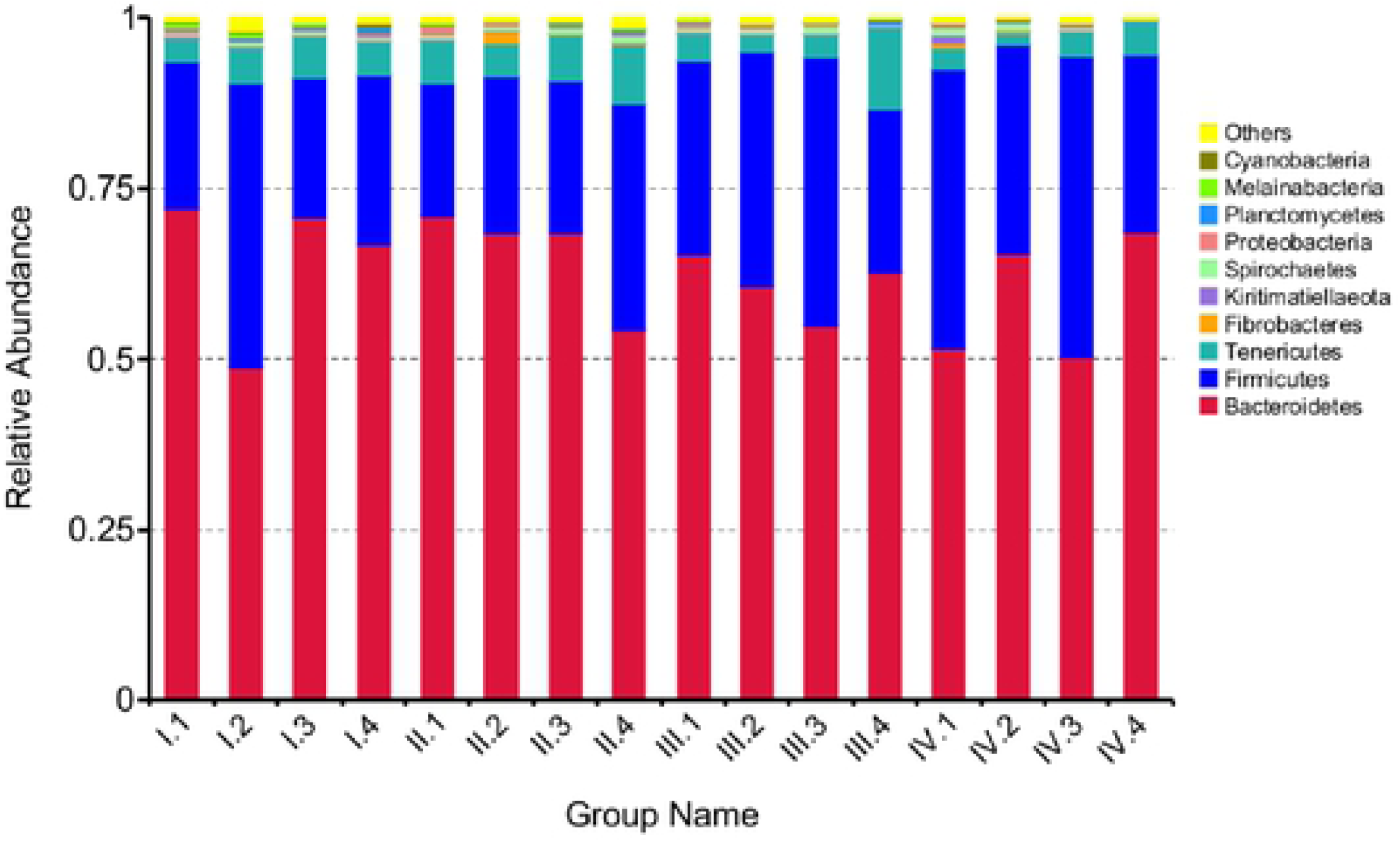
The column chart of the main dominant phylum in Karakul sheep fed with different NFC/NDF diets. A color-coded bar plot showing the average bacterial phylum distribution across the different age groups that were sampled.

**Fig. 5(a~b).**
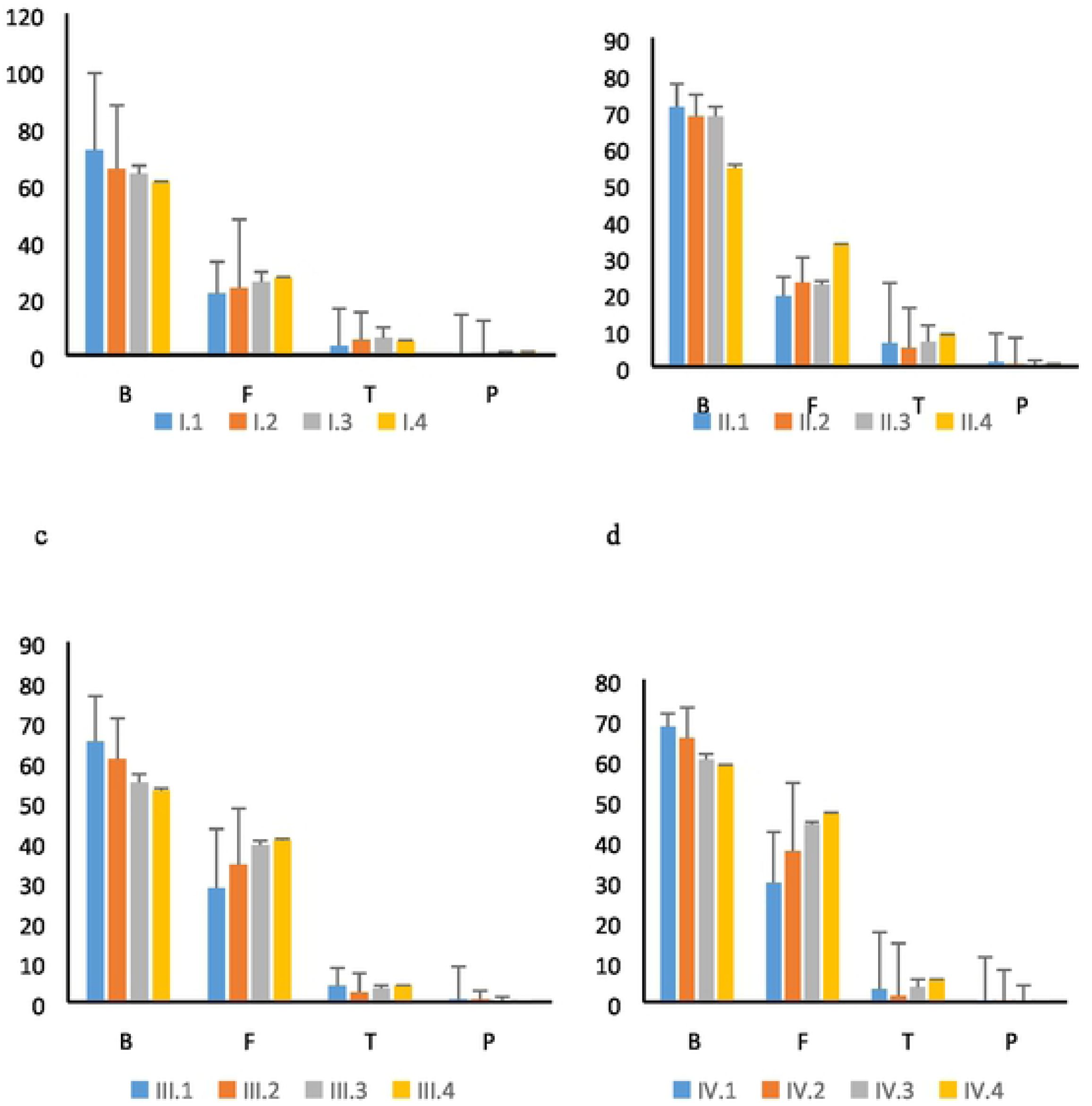
Effects of different NFC/NDF diets on relative abundance (% reads) of rumen phylum in Karakul Sheep. Note: B means Bactcroidcts, F means Firmicutcs, T means Tcncricutcs, P means Protcobactcria; a, b, c, d represents the experiment period ofl, II, III, IV respectively.

At the genus level, a total of 77 genera were obtained from sequence alignment. It can be seen from S3 Table and Fig. 6. The main genus microbiome didn’t change with the dietary NFC/NDF and periods. The highest relative abundance of genus was a kind of semi-cellulose degrading bacteria, *unidentified-Lachnospiraceae* (1.86%~16.68%). The following relative abundance of genus was *Succiniclasticum* (0.12%~17.03%). Fig. 7 was composed by four main dominant genus to investigate effects of dietary NFC/NDF on their relative abundance, the abundance of *Succiniclasticum* was that: group 2> group 1> group 3> group 4, and the difference wasn’t significant (*P*>0.05). The relative abundance of *unidentified-Lachnospiraceae*, *Anaeroplasma* and *unidentified-Bacteroidales* reached the highest in group 3 for four periods, and the difference wasn’t significant (*P*>0.05) as well.

**Fig. 6.**
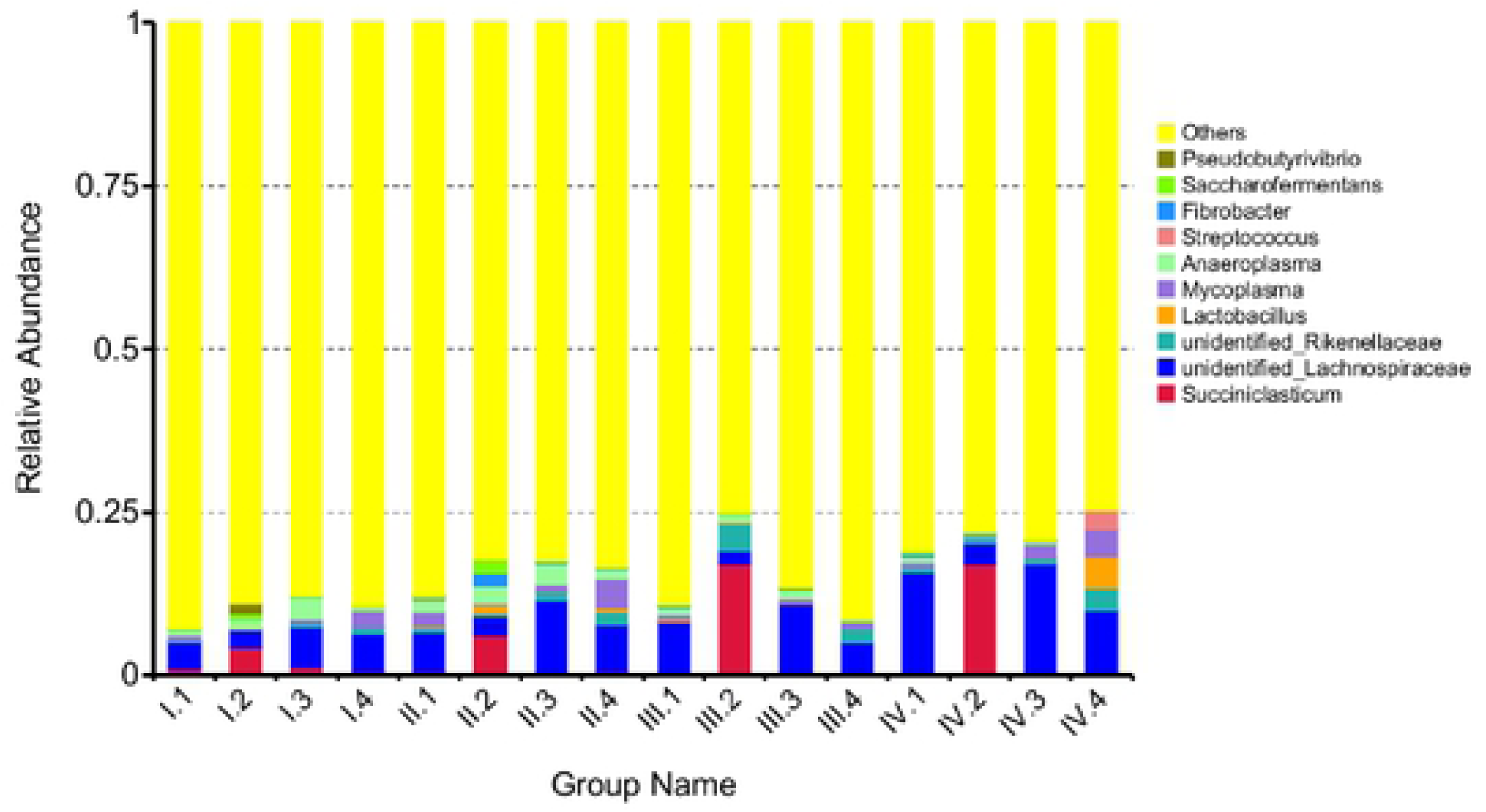
The column chart of the main dominant genus in Karakul sheep fed with different NFC/NDF diets. A color-coded bar plot showing the average bacterial genera distribution across the different age groups that were sampled

**Fig. 7 (a~d).**
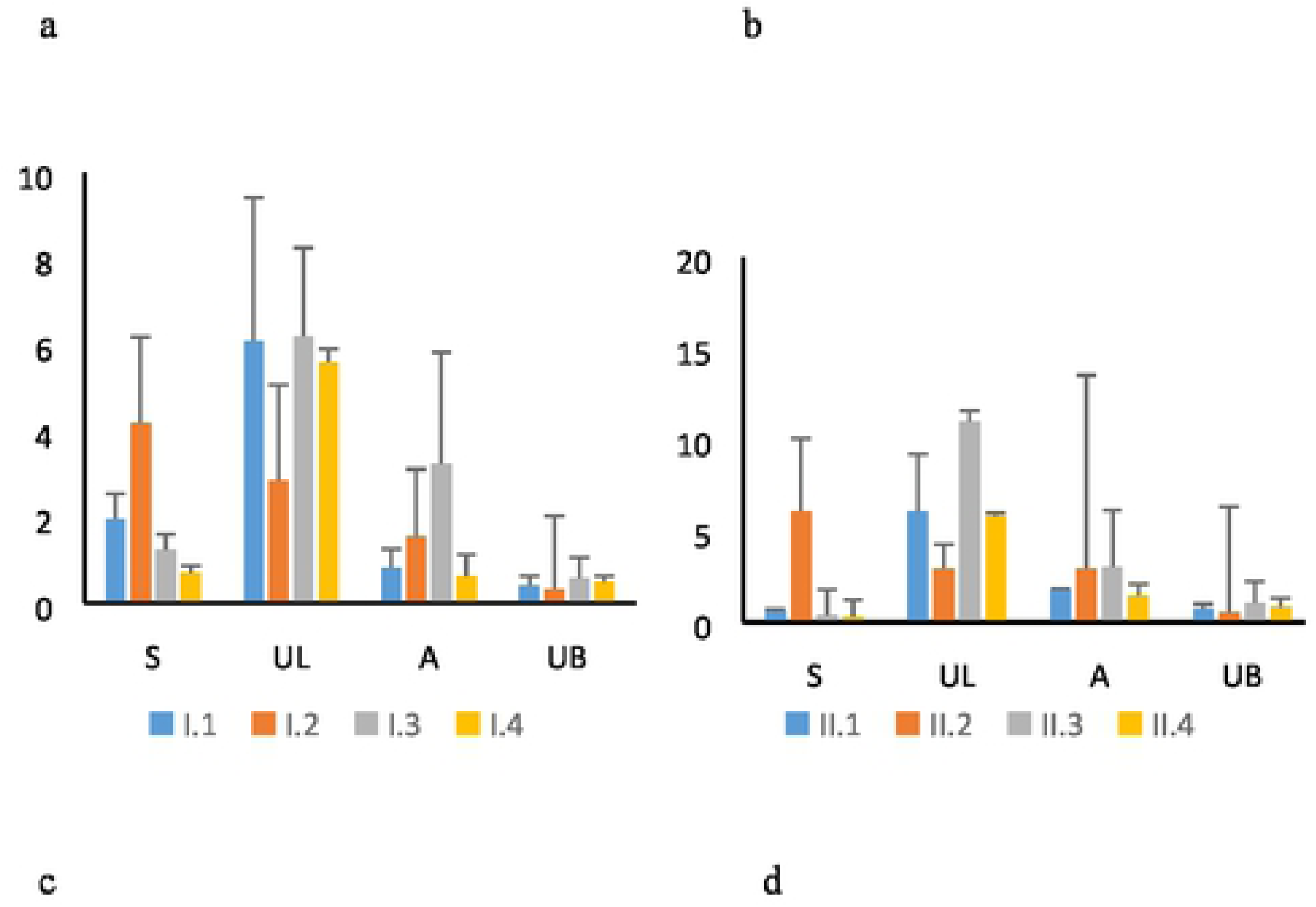
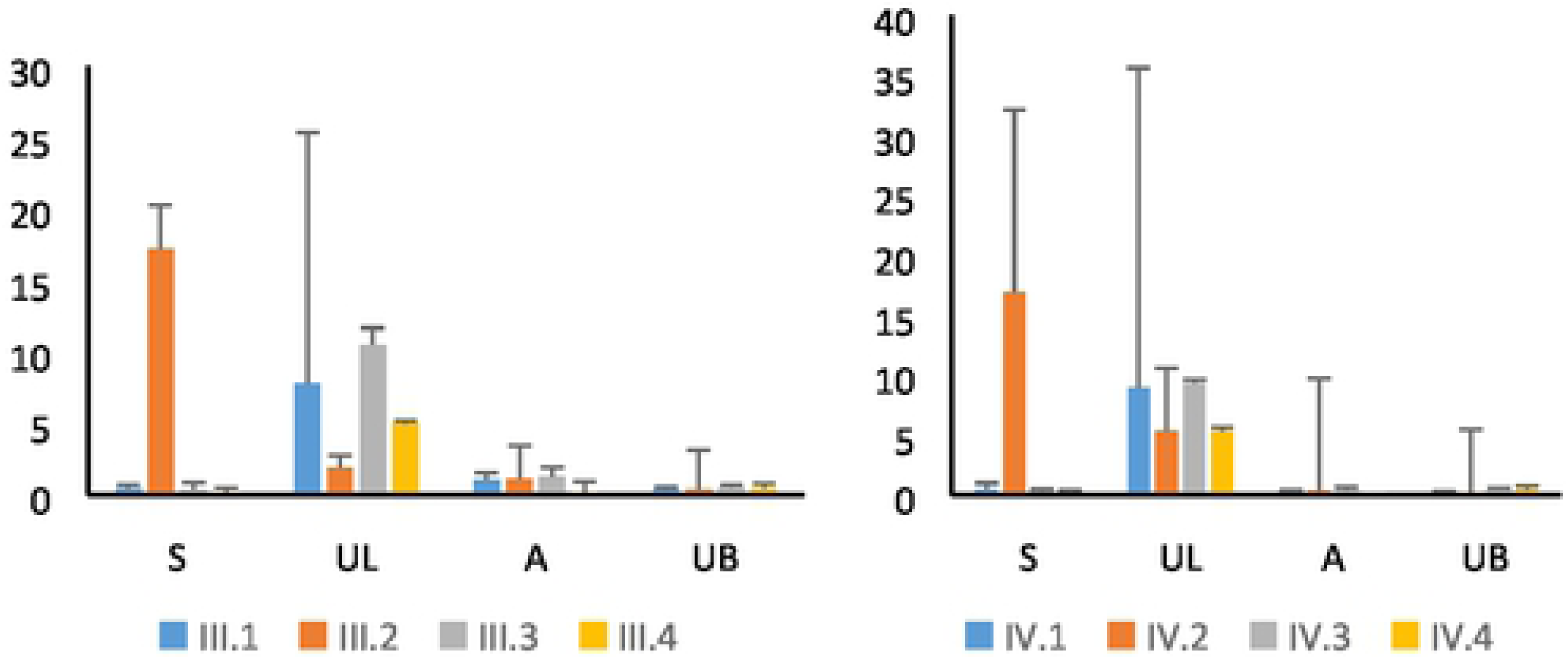
Effects of different NFC/NDF diets on relative abundance (% reads) of rumen genus in Karakul Sheep. Note: S means *Succiniclasticum*, UL means *unidentified-Lachnospiraceae*, A means *Anaeroplasma*, and UB means *unidentified Bacieroidales*, a, b, c, d represents the experiment period of I, II, III, IV respectively.

## Discussion

### Effects of dietary NFC/NDF on ruminal pH

pH is the directest index affecting rumen fermentation[25] and diets are key factors affecting pH. The results showed that the ruminal average pH decreased with the increase of NFC/NDF for four periods, which was approved with T.Ma et al.[26]. Agle et al. [27] and Pina et al. [28] also reported that with adding of concentration, ruminal pH decreased. This is mainly due to that the increase content of NFC resulted in the increase of VFA, while the low content of NDF resulted in the decreased rumination in sheep and decreased saliva to the rumen. Thus, the average pH of group IV was significantly lower than the other groups. The results were consistent with other studies[29–30]. Yang et al [31] pointed out that when pH was lower than 6.0 for a long time, the sheep would be in a long-term pathological state. The results showed that the average pH in group IV was near to 6.0, it may have negative effects on Karakul sheep, and needs to be further verified.

### Effects of dietary NFC/NDF on rumen bacteria in Karakul

#### sheep

Diets have crucial effects on rumen microorganisms, Jin et al. [32] showed that the number and diversity of rumen bacteria in goats feed under high grain (71.5%) diet were lower than those with high forage diet (0% grain). In this study, the results showed that the rumen bacteria diversity in Karakul sheep decreased with the increase of dietary NFC/NDF, which was consistent with the results of Liu [33], However Yong et al. [34] showed that there was no significant difference in the number of rumen bacteria when the sheep were fed with different ratio of forge to concentration diets, which might be caused by the difference of diets and species.

Roughage is the main feed source for ruminants, and rumen bacteria play a crucial role in the utilization of roughage. Research showed that diets with easily fermentable carbohydrates would decrease fiber degradation[35], resulting the imbalance of cellulolytic bacterial species. Bacteroidetes play an important role in the degradation of non-fiber substances and Firmicutes mainly degrade fiber substances. A large number of studies have shown that Bacteroidetes and Firmicutes are the most dominant flora in the gastrointestinal tract of mammals[36–39]. Li et al. [40] showed that when the calves were fed with two kinds of NFC/NDF diets, Bacteroidetes and Firmicutes were still the main dominant flora. In this study, the results showed that the relative abundance of Bacteroidetes and Firmicutes in different dietary NFC/NDF were still the main dominant phylum. Some results showed that the abundance of bacteria in the same sample would be different if the gene region was sequenced different[41], In our experiment, the region of V1-V9 was sequenced and the results showed that the relative abundance of rumen bacteroidetes decreased with the increase of dietary NFD/NDF in Karakul sheep, which was consistent with Ellison[42]. However, when Kim et al. [43] researched on the content of Bacteroidetes in beef cattle by sequencing V1-V3 region, the content of Bacteroidetes in high forage group was significantly lower than that of high proportion cereal group, which may be due to the difference of species differences and the sequence regions measured. In addition, the degradation rate of dry matter and organic matter was higher in group 3, 4 than which in group 1, 2 (the results were found previously by our team) so other bacteria except Bacteroidetes in Karakul sheep might have digested non-fiber substances and needs to be further studied.

Improving the fiber degradation rate is very important for ruminants. Bacteria and fungi play a crucial role in the decomposition and utilization of cellulose. In this study, the relative abundance of Firmicutes reached the highest when the dietary of NFC/NDF were 1.61 and 2.00, which was consistent with the results that had done before (The result was that: the NDF degradation rate in dietary NFC/NDF of 1.61 was the highest).U*nidentified-Lachnospiraceae* was the most dominant genus and its relative abundance reached the highest in group 3 for four periods, which further identified the results that: NDF degradation rate in dietary NFC/NDF of 1.61 was the highest.

In addition, there were many unidentified bacteria in the rumen of Karakul sheep, which might mean that there were some new species in Karakul sheep and needs to be further studied.

**Table 3.** Effects of different NFC/NDF on the relative abundance (%) of cellulose-degrading bacteria

| Period I |  |  |  |  |  |  |
| --- | --- | --- | --- | --- | --- | --- |
| Species | 1 | 2 | 3 | 4 | SEM | P-value |
| *Butyrivibrio-fibrisolvens* | 2.664a | 0.466b | 2.990a | 2.123a | 0.371 | 0.035 |
| *Fibrobacter-sp-UWCM* | - | 0.027 | - | - | 0.006 | 0.441 |
| *Ruminococcus-flavefaciens* | 0.027 | 0.054 | 0.051 | - | 0.038 | 0.216 |
| *Ruminococcus-albus* | - | 0.055 | - | - | 0.009 | 0.052 |
| Period II |  |  |  |  |  |  |
| Species | 1 | 2 | 3 | 4 | SEM | P-value |
| *Butyrivibrio-fibrisolvens* | 2.689a | 0.548b | 3.123a | 2.406a | 1.132 | 0.027 |
| *Fibrobacter-sp-UWCM* | 0.082 | 0.411 | 0.027 | - | 0.350 | 0.294 |
| *Ruminococcus-flavefaciens* | - | 0.055 | 0.050 | 0.027 | 0.036 | 0.290 |
| *Ruminococcus-albus* | 0.027 | 0.055 | - | - | 0.01 | 0.561 |
| Period III |  |  |  |  |  |  |
| Species | 1 | 2 | 3 | 4 | SEM | P-value |
| *Butyrivibrio-fibrisolvens* | 6.785a | 0.657c | 7.687a | 3.292b | 1.143 | <0.01 |
| *Fibrobacter-sp-UWCM* | 0.025 | 0.027 | - | - | 0.009 | 0.596 |
| *Ruminococcus-flavefaciens* | 0.027 | 0.082 | 0.055 | 0.027 | 0.021 | 0.627 |
| *Ruminococcus-albus* | - | - | - | - | - | - |

Period IV
| Species | 1 | 2 | 3 | 4 | SEM | P-value |
| --- | --- | --- | --- | --- | --- | --- |
| *Butyrivibrio-fibrisolvens* | 7.234a | 2.301c | 8.694a | 5.975b | 1.581 | <0.01 |
| *Fibrobacter-sp-UWCM* | - | 0.082 | - | - | 0.07 | 0.441 |
| *Ruminococcus-flavefaciens* | - | 0.082 | 0.079 | - | 0.016 | 0.052 |
| *Ruminococcus-albus* | - | 0.085 | - | - | 0.013 | 0.063 |

**Fig. 8.**
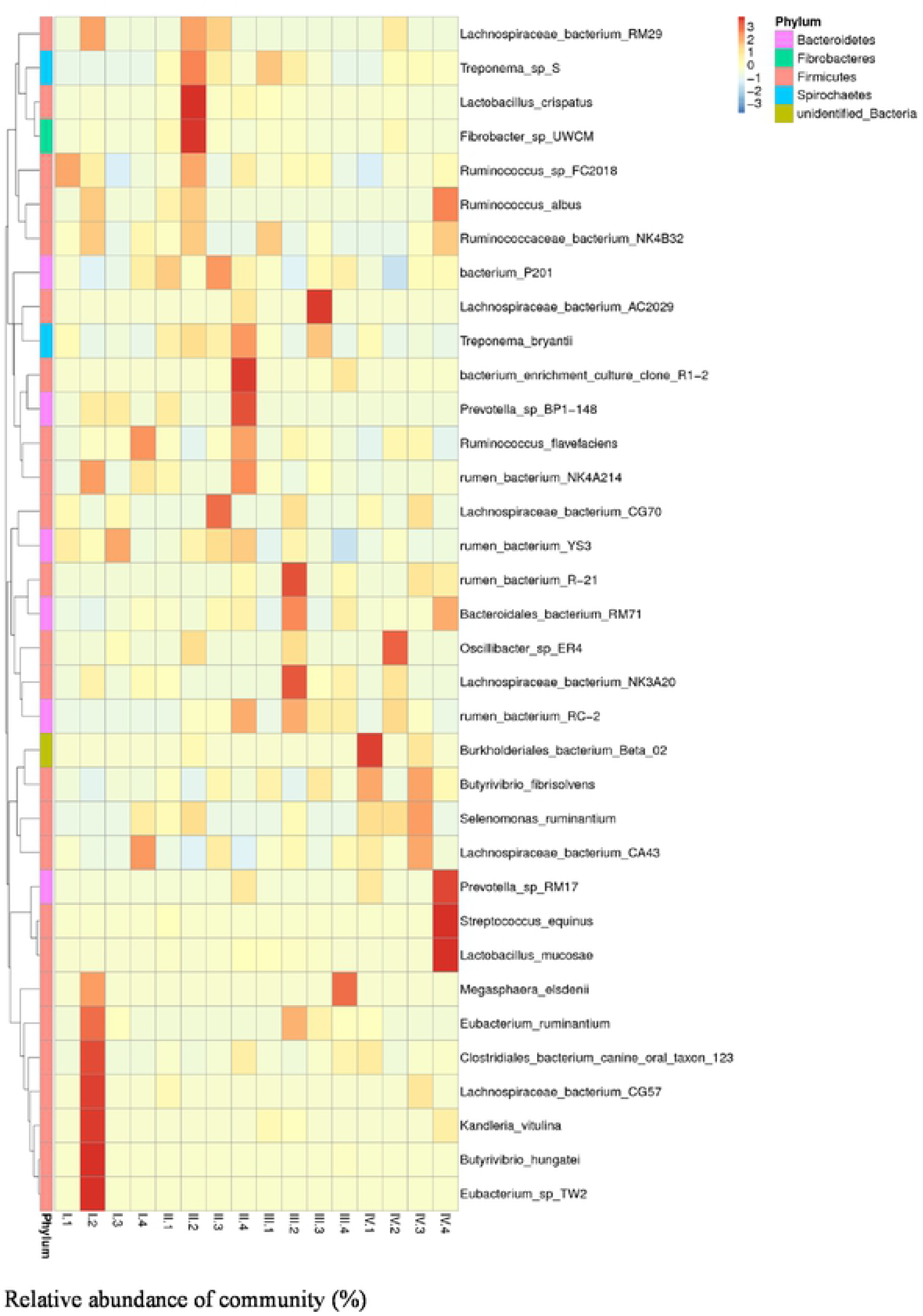
Heat mop of the rumen bacteria composition at species level. The heat map indicates the relative percentage of each species for the different dietary NFC/NDF group sampled.

## Conclusions

The ruminal pH and total diversity of rumen bacteria decreased with the increase of dietary NFC/NDF. The most dominant phylum, genus and species didn’t change with dietary NFC/NDF and the ruminal bacteria became more stable with prolong of periods in Karakul sheep.

## Acknowledgments

We are grateful for other tutors and classmates in our department that helped in the experiment.

## Author contributions

Contributed reagents/materials/analysis/ tools: X.G.; Performed the experiments: X.P; Analyzed the data: X.P; Writing-original draft: X.P.; Writing–review editing: X.P., X.G., C.J., J.L., X.Z., S.Z., C.L., and A.S..

## Supporting information

**S1 Table. The ingredients and nutrient composition of the diet (% of DM).** ①The premix provided the following per kg of diets: VA 1800 IU, VD3 600 IU, VE 30 mg, Fe 65 mg, Se 0.15 mg, I 0.6 mg, Cu 10 mg, Mn 28 mg, Zn 45 mg, Cu 12 mg. ②Nutrition level was a calculated value. ③NFC=(1NDFCPFatAsh)× 100%。

**S2 Table Effects of dietary NFC/NDF on relative abundance of phylum in Karakul sheep**

Period I (1~18 d), II (19~36 d), III (37~54 d) and IV (55~72 d) and Group 1, 2, 3, 4 means four groups of sheep treated with four dietary levels of NFC/NDF (0.78, 1.23, 1.61, 2.00 respectively). The same as below.

**S3 Table Effects of dietary NFC/NDF on relative abundance of genus in Karakul sheep**

